# Adult Neurogenesis Conserves Hippocampal Learning Capacity

**DOI:** 10.1101/200253

**Authors:** Md Jahangir Alam, Takashi Kitamura, Yoshito Saitoh, Noriaki Ohkawa, Takashi Kondo, Kaoru Inokuchi

## Abstract

Memory coding strengthens synaptic efficacy in the hippocampus via a long-term potentiation (LTP)-like mechanism. Given that animals are able to store memories of everyday experiences, hippocampal circuits should be able to avoid saturation of overall synaptic weight to preserve learning capacity. However, the underlying mechanism for this is still poorly understood. Here, we show that adult neurogenesis in rats plays a crucial role in the maintenance of the hippocampal learning capacity for learning. Artificial saturation with hippocampal LTP impaired learning capacity in contextual fear conditioning, which then completely recovered after 14 days, when LTP had decayed to the basal level. Ablation of neurogenesis by X-ray irradiation delayed the recovery of learning capacity, while enhancement of neurogenesis using running wheel sped up the recovery. Thus, one benefit of ongoing adult neurogenesis is the maintenance of hippocampal learning capacity through homeostatic renewing of hippocampal memory circuits. Decreased neurogenesis in aged animals may underlie declines in cognitive function with aging.

## INTRODUCTION

The hippocampus is crucial for declarative memories in humans (Ofen et al., 2007; Scoville and Milner, 1957), and encodes episodic and spatial memories in animals (Buzsaki and Moser, 2013; Conway, 2009). During memory formation, synaptic plasticity at appropriate synapses is both necessary and sufficient for information storage (Nabavi et al., 2014). Hippocampus-dependent learning basically induces strengthening of synaptic efficacy through long-term potentiation (LTP) in the hippocampus (Hebb, 1949; Tononi and Cirelli, 2014; Whitlock et al., 2006), which implies that information coding leads to the saturation of neuronal circuits. Sleep plays an important role in memory formation, by removing and down-sizing spines, and thereby scaling down excitatory synapses (de Vivo et al., 2017; Li et al., 2017; Tononi and Cirelli, 2014) to avoid saturation of the circuits. However, accumulation of memories selected for long-lasting storage would eventually lead to a net increase in synaptic strength and saturate the neural circuits, which in turn would lead to the hippocampus being unable to store new memories. Thus, a mechanism other than sleep-mediated scaling down may work to prevent the saturation of hippocampal circuits.

Throughout adulthood, new neurons are continuously generated and functionally integrated into the existing circuits in the hippocampal dentate gyrus (DG) (Altman and Das, 1965; Deng et al., 2010; Kitamura and Inokuchi, 2014; Zhao et al., 2008). Hippocampal neurogenesis modulates the hippocampus-dependent period of contextual fear memory, where inhibition or enhancement of hippocampal neurogenesis respectively prolongs or shortens the hippocampus-dependent period of memory (Kitamura and Inokuchi, 2014; Kitamura et al., 2009). However, little is known how brain avoids the saturation state and retains the learning capacity. In the present study, we show that adult neurogenesis plays an important role in the maintenance of hippocampal learning capacity and enables animals to learn new events through the renewal of hippocampal memory circuit.

## RESULTS

### Recovery of Learning Capacity from Saturation level Correlates with LTP Decay

Two experimental approaches were employed to saturate hippocampal synaptic efficacy in rats: repeated high-frequency tetanic stimulation (rHFS) (Castro et al., 1989; Moser et al., 1998) and repeated maximum electroconvulsive shock (rMECS) (Stewart et al., 1994). In the first series of experiments, hippocampal synaptic efficacy was saturated by delivering rHFS to the perforant pathway (PP) of the right hemisphere, which projects to the DG of the dorsal hippocampus. The dorsal hippocampus of the left hemisphere was damaged by injection of ibotenic acid (IBO) into multiple sites(Jarrard, 1989) (Table S1). Cresyl violet staining showed that the multiple IBO injections completely damaged the dorsal hippocampus (Figure S1A and S1B). This unilateral IBO injection had no effect on hippocampus-dependent learning, contextual fear conditioning (CFC), in comparison with its vehicle (phosphate-buffered saline; PBS) injection control (Figure S1C). After a 10 day period for recovery from the IBO injection, stimulating and recording electrodes were implanted into the right hemisphere (Figure 1A and 1B). rHFS was delivered to the PP-DG pathway, and the evoked synaptic responses were measured from the DG in freely moving non-anesthetized rats. The application of rHFS gradually increased the slope of the field excitatory postsynaptic potentiation (fEPSP slope), which reached saturation level after the fifth stimulation session (Figure 1C and 1D). One day after the last HFS session, the animals were trained with the CFC task. The rHFS group showed lower freezing in the test session 1 day after the training when compared with the test pulse group (Figure 1E). DG-LTP saturation had no effect on auditory fear conditioning (AFC), a hippocampus-independent learning task, as both the rHFS and test pulse groups showed the same freezing in the tone session (Figure 1F and 1G). Thus, saturation of hippocampal LTP impaired hippocampus-dependent learning, without affecting hippocampus-independent learning.

**Figure 1.**
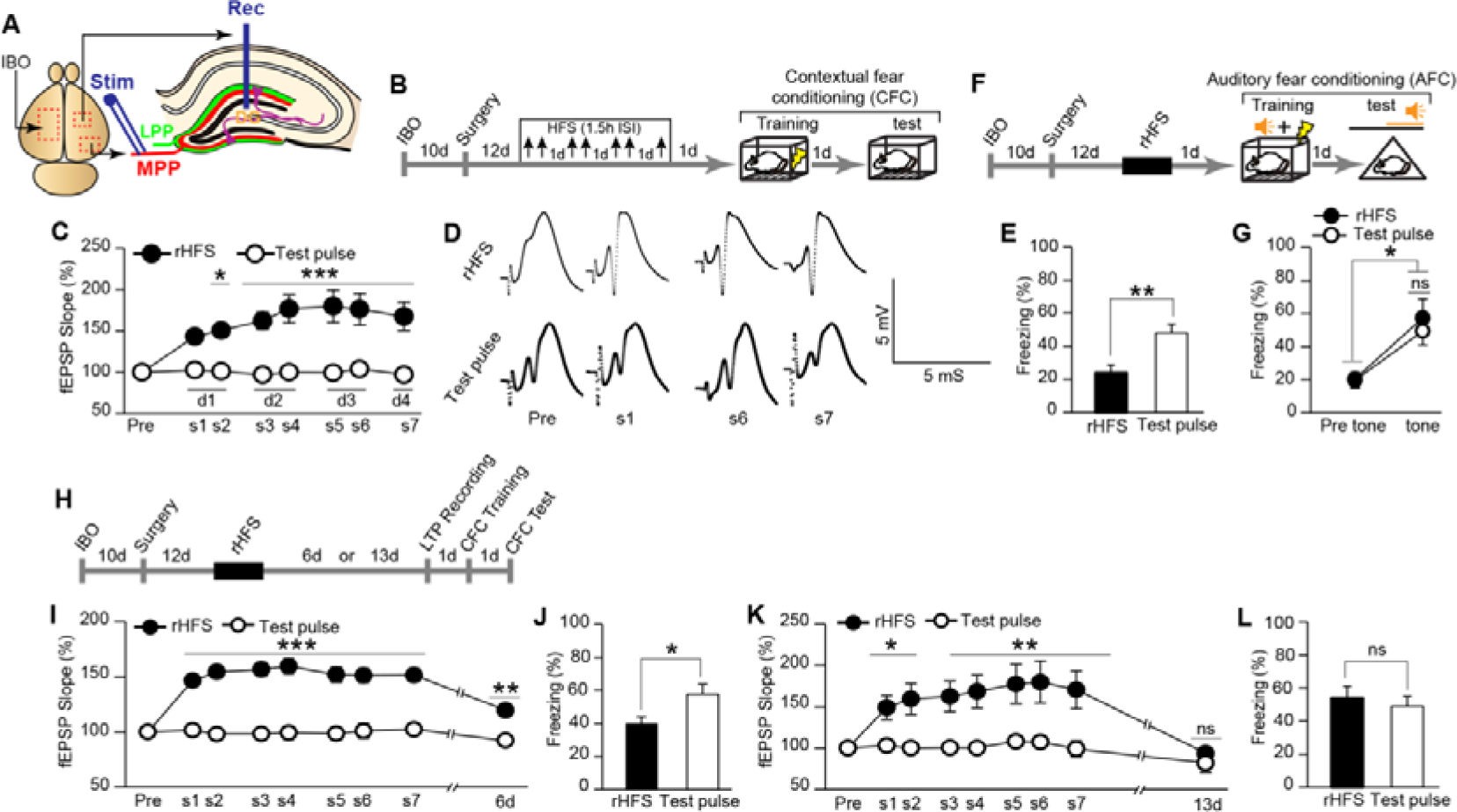
Recovery from hippocampal LTP saturation correlates with the learning capacity for CFC. (A) Schematic illustration showing experimental set up for IBO injection and electrode implantation. MPP, medial perforant path; LPP, lateral perforant path; Rec, recording electrode; Stim, stimulating electrode; IBO, ibotenic acid.(B) Experimental schedule for the effect of rHFS. Upper arrows indicate the stimulation session.(C) Saturation of hippocampal LTP. The normalized values for the fEPSP slope for rHFS (*n* = 9) and Test pulse (*n* = 7) rats (two-way ANOVA, *P* = 0.0009, *F*_1,14_ = 17.58, Bonferroni corrected for multiple comparisons). s, session; d, day.(D) Example traces of evoked fEPSP responses. Pre, before rHFS.(E) Average freezing response during the test session (rHFS, *n* = 9; Test pulse, *n* = 7 rats; unpaired *t*-test, *P* = 0.0051, *t*_14_ = 3.31).(F) Experimental schedule for AFC.(G) Freezing response before tone (pre) and during tone period (tone; *n* = 9 rats/group) in the AFC test (paired *t*-test, rHFS pre-tone vs tone, *P* = 0.01, *t*_8_ = 3.18; Test pulse pre-tone vs tone, *P* = 0.01, *t*_8_ = 3.19; unpaired *t*-test, rHFS tone vs Test pulse tone, *P* = 0.58, *t*_16_ = 0.56).(H) Experimental schedule for the gradual recovery of synaptic efficacy and learning capacity.(I) LTP partially decayed after 6 days (two-way ANOVA, *P* < 0.0001, *F*_1,15_ = 63.82; rHFS d6 vs Test pulse d6, *P* = 0.0012, Bonferroni corrected for multiple comparisons). (J) Incomplete recovery of learning capacity (unpaired *t*-test, *P* = 0.03, *t*_15_ = 2.29) 7 days after the last HFS (Test pulse, *n* = 9; rHFS, *n* = 8 rats).(K) LTP returned to the baseline level 13 days after rHFS (two-way ANOVA, *P* = 0.0065, *F*_1,14_ = 10.18; rHFS d13 vs Test pulse day 13, *P* > 0.99, Bonferroni corrected for multiple comparisons).(L) Recovery of learning capacity (unpaired *t*-test, *P* = 0.60, *t*_14_ = 0.53, *n* = 8 rats/group). * *P* < 0.05, ** *P* < 0.01, *** *P* < 0.001; ns, not significant.

LTP was shown to have partially decayed from the saturation level 6 days after the last HFS session (Figure 1H and 1I). Next day the animals were trained with the CFC task. The rHFS group showed lower freezing than the test pulse group during the test session on the following day (Figure 1H and 1J), but higher freezing than the rHFS group, in which CFC training was performed 1 day after the rHFS (see Figure 1E). LTP had completely decayed with a return to the baseline level 13 days after the last HFS (Figure 1H and 1K). The following day, the animals were trained with the CFC task. Both test pulse and rHFS groups showed comparable freezing in the test session (Figure 1L). General activity during the training session of the CFC task was comparable between the rHFS and control groups (Figure S2A to S2C). Thus, learning capacity gradually recovered from saturation in parallel with the gradual decay of LTP.

### Decreased Hippocampal Neurogenesis Extends the Recovery of Learning Capacity

We then assessed the role played by neurogenesis in the recovery of learning capacity. Rats were injected with IBO, and the whole brain was focally irradiated 10 days later, followed by LTP saturation and CFC training (Figure 2A). rHFS in both the non-irradiated and irradiated groups saturated the hippocampal LTP, with no significant between-group difference in the magnitude of LTP (Figure 2B and 2C). LTP in the non-irradiated group had decayed and returned to the basal level 13 days after the last HFS. By contrast, the irradiated group retained the LTP for 13 days. On the following day, the animals were trained with the CFC task. The irradiated group showed significantly lower freezing in test sessions when compared with the non-irradiated group (Figure 2D), and also had significantly reduced neurogenesis in comparison with the non-irradiated group (Figure. 2E to 2G). Both groups showed comparable motility in the training session (Figure S2D). X-ray irradiation alone had no effect on the CFC learning ability, as irradiated rats showed freezing in the test session 1 day after the CFC training comparable with that shown by the non-irradiated rats (Figure 2H and 2I). Motility during the training session was not affected by irradiation (Figure S2E). There was a significant negative correlation between the fEPSP slope and freezing (Figure 2J). Thus, the reduced neurogenesis postponed the recovery of hippocampus-dependent learning capacity by maintaining the saturated level of hippocampal LTP.

**Figure 2.**
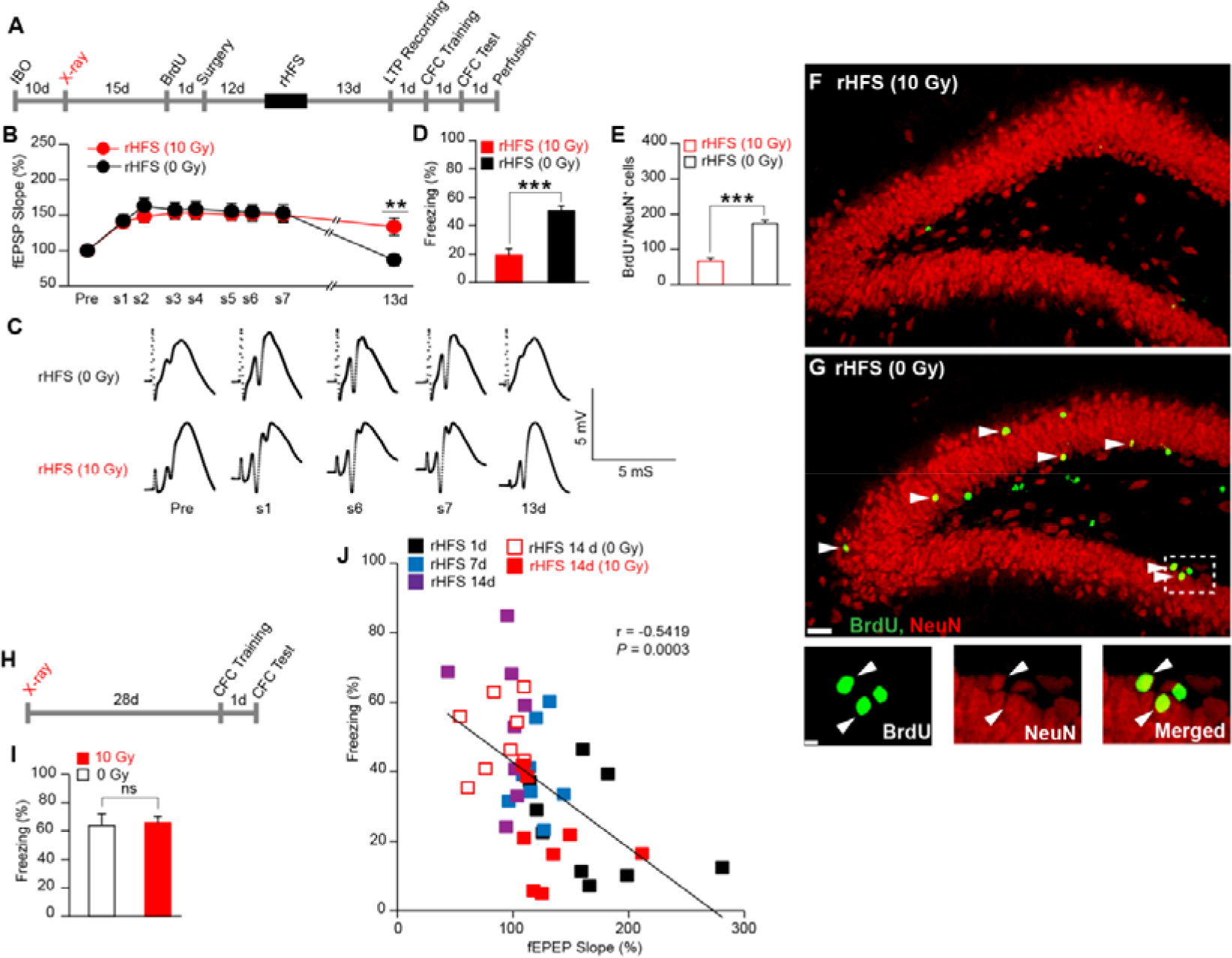
Adult hippocampal neurogenesis regulates the recovery from rHFS-induced impairment of CFC. (A) Experimental schedule.(B) LTP maintenance in non-irradiated (0 Gy) and X-ray-irradiated (10 Gy) rats [two-way ANOVA, *P* = 0.91, *F*_1,14_ = 0.01; rHFS (10 Gy) 13day vs rHFS (0 Gy) 13day, unpaired *t*-test, *P* = 0.005, *t*_14_ = 3.25, *n* = 8 rats/group].(C) Example traces of evoked fEPSP responses. Pre, before rHFS.(D) Freezing response during the test session (unpaired *t*-test, *P* = 0.0002, *t*_14_ = 4.87, *n* = 8 rats/group). (E) Reduced neurogenesis in the rHFS-irradiated rats (n = 3 rats/group, unpaired *t*-test, *P* = 0.0009, *t*_4_ = 8.77).(F and G) Hippocampal coronal sections. Scale bar, 200 μm. Arrowheads indicate BrdU^+^/NeuN^+^ double positive cells. (G) Lower panel, magnified images from rHFS (0 Gy): BrdU (left), NeuN (middle) and merged (right). Scale bar, 50 μm.(H) Experimental schedule for the effect of X-rays on CFC.(I) Freezing response at test (10 Gy, *n* = 8; 0 Gy, *n* = 7 rats; unpaired *t*-test, *P* = 0.84, *t*_13_ = 0.19).(J) Correlation of LTP and learning ability (Pearson correlation test, *r* = −0.5419, *P* = 0.0003). Each dot represents data from an individual animal. *P < 0.05, \*\**P* < 0.01, \*\*\**P* < 0.001; ns, not significant. *, runner vs irradiated runner; runner vs non-runner; †, non-runner vs irradiated runner.

### Increased Hippocampal Neurogenesis Sped up the Recovery of Learning Capacity

Voluntary running wheel exercise (Kitamura et al., 2009; van Praag et al., 1999) was employed to examine the effect of increased neurogenesis on the recovery from saturation level (Figure 3). From 1 day after the irradiation, rats were housed in running wheel cages for the entire remaining session, except for the 5 days immediately after the IBO injection (Figure 3A). The distance run was comparable between the non-irradiated and irradiated rats (Figure 3B). One month after the onset of the exercise regimen, rats were subjected to rHFS. There was no between-group difference in the fEPSP slope during the rHFS (Figure 3C and 3D). When measured 6 days after the rHFS; runner rats showed faster LTP decay from saturation level than the non-runner and irradiated runner groups (Figure 3C). The next day, the animals were subjected to CFC training. The runner group showed complete recovery of learning capacity, whereas the other groups showed impaired learning capacity (Figure 3E and 3F; see also Figure 1J). Rats were administered BrdU 1 day after the CFC test, and 2 h later they were perfused and analyzed for cell proliferation. The 6 weeks of running exercise enhanced cell proliferation in the subgranular zone of the DG in the runner group in comparison with the non-runner and irradiated runner groups (Figure 3G to 3J). Data from individual animals showed a significant negative correlation between fEPSP slope and freezing (Figure 3K). Taken together, ablation or enhancement of neurogenesis respectively delayed or accelerated LTP decay, which was accompanied by slow or rapid recovery of learning capacity (Figure 3L and 3M).

**Figure 3.**
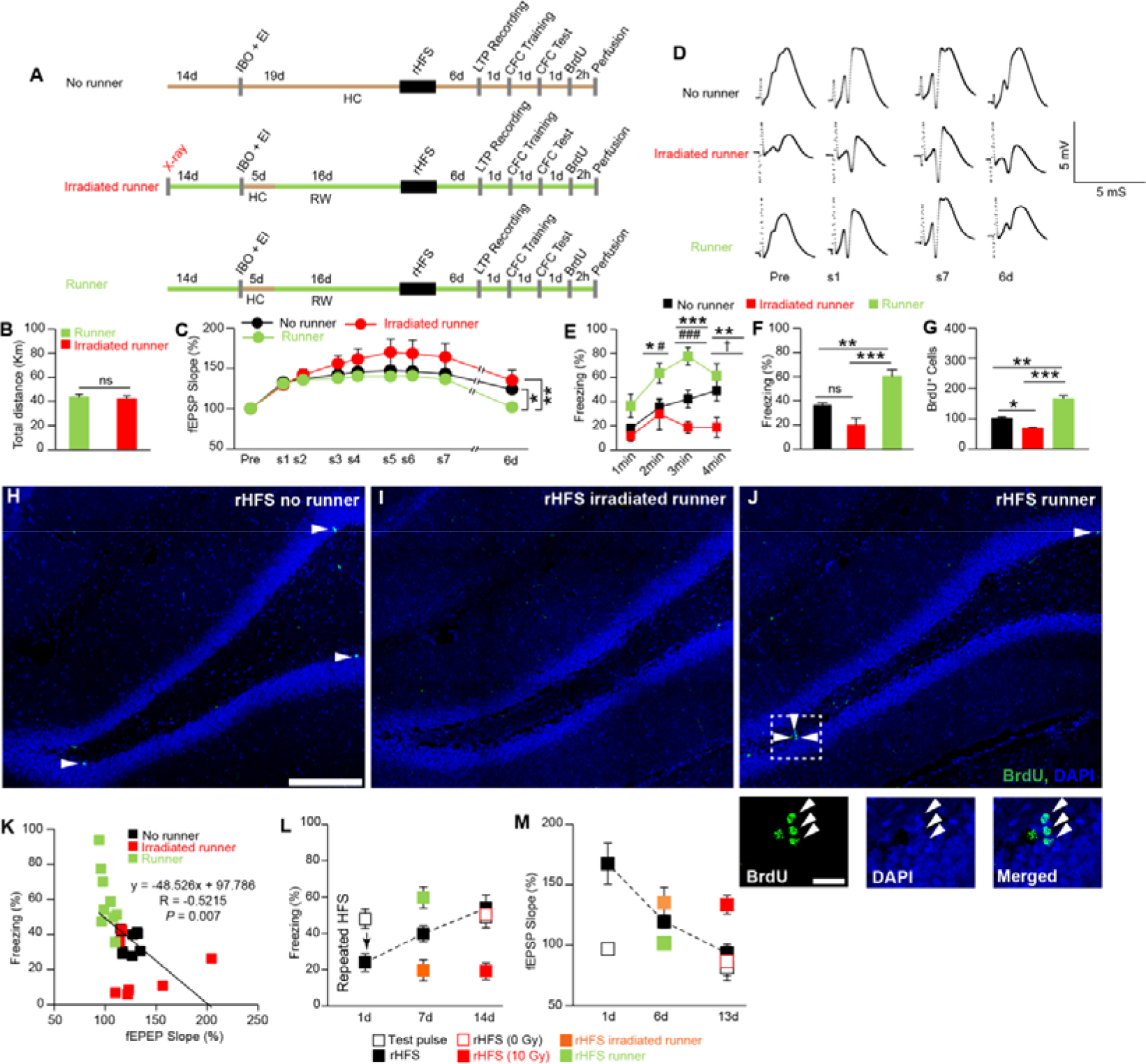
Adult hippocampal neurogenesis accelerates the recovery of rHFS-induced learning impairment. (A) Experimental schedule. Green or ochre bars indicate housed with (running wheel, RW) or without (home cage, HC) a running wheel, respectively. IBO + EI, ibotenic acid injection and electrode implantation.(B) Total distance travelled (runner, *n* = 9; irradiated runner, *n* = 7 rats; unpaired *t*-test, *P* = 0.72, *t*_14_ = 0.36).(C) LTP maintenance [two-way ANOVA, *P* = 0.07, *F*_2,22_ = 2.89 (one-way ANOVA, *P* = 0.005, *F*_2,22_ = 6.54, Fisher’s LSD post-hoc test, runner 6d vs non-runner 6d, *P* = 0.02; runner 6d vs irradiated runner 6d, *P* = 0.002), non-runner, *n* = 9; runner, *n* = 9; irradiated runner, *n* = 7 rats].(D) Example traces of evoked fEPSP responses. Pre, before rHFS.(E and F) Time courses of freezing (E, two-way ANOVA, *P* < 0.0001, *F*_2,22_ = 17.04, Tukey’s multiple comparisons test) and average freezing responses (F, one-way ANOVA, *P* < 0.0001, *F*_2,22_ = 17.0, Tukey’s multiple comparisons test) during CFC test (non-runner, *n* = 9; runner, *n* = 9; irradiated runner, *n* = 7 rats).(G) Proliferation of the new neurons (*n* = 3 rats/group, one-way ANOVA, *P* = 0.0001, *F*_2,6_ = 58.1, Tukey’s multiple comparisons test).(H to J) Hippocampal coronal sections from a non-runner (H), an irradiated runner (I) and a runner rats (J). Scale bar, 100 μm. Arrowheads indicate BrdU^+^ positive cells. (J) Lower panels, magnified images. Scale bar, 50 μm. BrdU (left), DAPI (middle) and merged (right).(K) Correlation of hippocampal LTP and learning ability (Pearson correlation test, *r* = −0.5215, *P* = 0.0075). Each dot represents data from an individual animal.(L and M) Summary of the behavioral (L) and LTP (M) results from Figures 1-3. * *P* < 0.05, ** *P* < 0.01, *** *P* < 0.001; ns, not significant. # *P* < 0.05, ### *P* < 0.001 (runner vs non-runner), † *P* < 0.05 (non-runner vs irradiated runner).

### Adult Hippocampal Neurogenesis Assists the Recovery from rMECS-induced Learning Impairment

Whole brain stimulation with rMECS is another way to saturate hippocampal synaptic efficacy(Stewart et al., 1994). Rats received 10 MECS and were then trained with the CFC task (Figure 4A). The rMECS-2day group, in which the interval between the last MECS and CFC training was 2 days, showed lower freezing than the sham group in the CFC test session 1 day after the training (Figure 4B). LTP induced after the CFC was impaired in the rMECS group in comparison with the no rMECS sham group (Figure 4C and 4D). rMECS had no effect on the hippocampus-independent learning task (Figure 4E and 4F). Thus, as was the case for rHFS, rMECS saturated hippocampal LTP and impaired hippocampus-dependent learning capacity.

**Figure 4.**
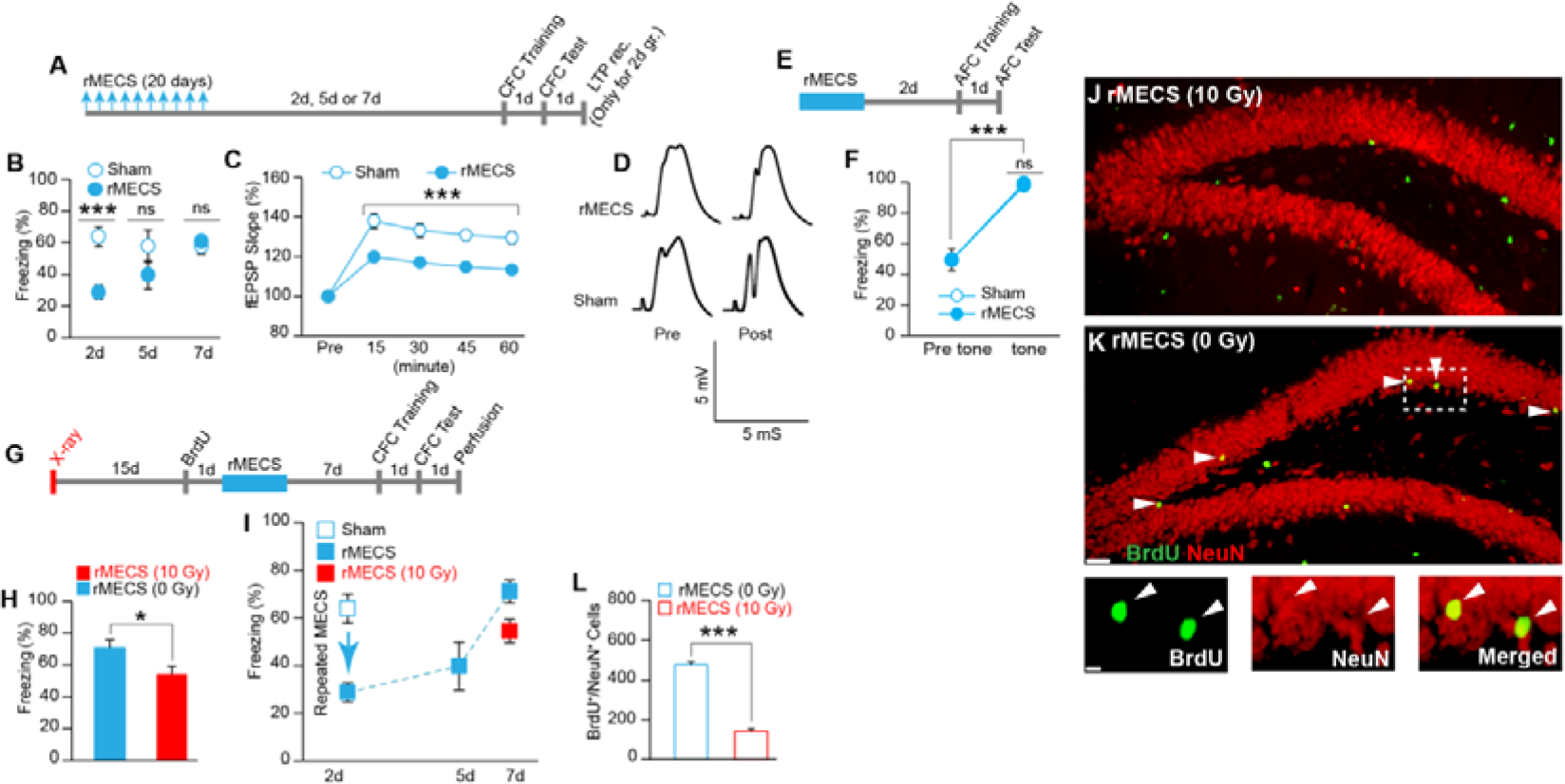
Adult hippocampal neurogenesis facilitates recovery from rMECS-induced impairment of CFC. (A) Experimental schedule. Upper arrows indicate the MECS induction.(B) Mean freezing response during the test. rMECS-2d groups (unpaired *t*-test, *P* < 0.0001, *t*_24_ = 4.72, *n* = 13 rats/group), 5d groups (unpaired *t*-test, *P* = 0.20, *t*_11_ = 1.33, rMECS, *n* = 7; Sham, *n* = 6 rats) or 7d groups (unpaired *t*-test, *P* = 0.58, *t*_23_ = 0.55, rMECS, *n* = 15; Sham, *n* = 10 rats) after the last MECS.(C) LTP induction in PP-DG synapses after CFC (two-way ANOVA, *P* = 0.0002, *F*_1,15_ = 25.04, Bonferroni corrected, Sham, *n* = 7; rMECS, *n* = 10 rats).(D) Example traces of evoked fEPSP responses recorded before (pre) and after (post) tetanization.(E) Experimental schedule for AFC.(F) Averaged freezing response before tone (pre) and during tone period (tone; Wilcoxon matched-pairs signed rank test; rMECS pre-tone vs tone, *P* = 0.003, Sham pre-tone vs tone, *P* = 0.002; rMECS tone vs Sham tone, unpaired *t*-test, *P* = 0.58, *t*_16_ = 0.56; Sham, *n* = 10; rMECS, *n* = 9 rats).(G) Experimental schedule for CFC.(H) Freezing response during the CFC test (unpaired *t*-test, *P* = 0.02, *t*_18_ = 2.41, *n* = 10 rats/group).(I) Summary of the behavioral results.(J and K) Hippocampal coronal section from rMECS-irradiated (J) and rMECS-non-irradiated rats (K). Scale bar, 200 μm. Arrowheads indicate BrdU^+^/NeuN^+^ double positive cells. Magnified images from an rMECS-non-irradiated rat (K; lower panel); BrdU (left), NeuN (middle) and merged (right). Scale bar, 50 μm.(L) Number of BrdU^+^/NeuN^+^ double positive cells (*n* = 4 rats/group, unpaired *t*-test, *P* < 0.0001, *t*_6_ = 16.99). * *P* < 0.05, ** *P* < 0.01, *** *P* < 0.001; ns, not significant.

The rMECS-7day group showed comparable freezing to the sham group in the CFC test session (Figure 4B). Thus, the saturation of learning capacity caused by rMECS had completely recovered after 7 days. The motility of the animals during the training session of the CFC task was comparable between the rMECS and sham groups (Figure. S3A to S3D), indicating that rMECS had no effect on the general activity of the animals.

The effect of irradiation was assessed by subjecting animals to the same course of rMECS 18 days after irradiation (Figure 4G). Animals were trained with the CFC task 7 days after the last MECS. In the test session 1 day after the CFC training, the X-ray-irradiated group (rMECS + 10 Gy) showed lower freezing than the non-irradiated group (rMECS + 0 Gy; Fig. 4G to 4I). The irradiated group showed reduced neurogenesis in comparison with the non-irradiated group (Figure 4J to 4L).

## DISCUSSION

This study demonstrates the existence of a circuit mechanism for homeostatic plasticity, which is mediated by adult neurogenesis to avoid saturation of hippocampal circuits. Hippocampal neurogenesis slowly renormalizes the enhanced synaptic strength resulting from LTP induced by strong tetanic stimulation, with the return to basal level usually taking around 2 weeks. Thus, sleep (de Vivo et al., 2017; Li et al., 2017; Tononi and Cirelli, 2014) and neurogenesis may play complementary roles by respectively mediating rapid and slow scaling down of synaptic strength. As hippocampal neurogenesis specifically renormalizes the strong tetanus-induced long-lasting LTP, it may result in the decay of consolidated memories, which escape the sleep-mediated erasure from the hippocampus. This neurogenesis-mediated decay of hippocampal memory does not result in the loss of the memory, as this process is accompanied by an increase in the neocortex dependence of the memory (Kitamura et al., 2009).

Synaptic competition between old and new neurons occurs when new neurons are integrated into pre-existing circuits (Lledo et al., 2006; Toni et al., 2007; Zhao et al., 2008). Integration of new neurons has an impact on activity-dependent synaptic rewiring in the hippocampus (Ohkawa et al., 2012; Toni et al., 2007). Thus, a blockade of new synapse integration stabilizes the hippocampal network, which then prevents the clearance of memories from the hippocampus. The replacement of synapses in this manner may maintain learning capacity, because the naive synaptic connections made by the new neurons are flexible, and can, therefore, be integrated into various networks that store newly acquired memories.

The level of adult hippocampal neurogenesis is regulated by many factors (Deng et al., 2010; Gould et al., 1999; Kitamura and Inokuchi, 2014; Ohkawa et al., 2012; Zhao et al., 2008) including neuronal activity, stress, and aging. A decrease in neurogenesis with aging is associated with an age-related decline in some forms of hippocampus-dependent memories (van Praag et al., 2005; Villeda et al., 2011). Given the finite storage capacity of the hippocampus, reduced neurogenesis in aged animals may underlie the limited capacity of the hippocampus to acquire and store new information, by reducing the clearance of old memories that have already been stored in cortical networks.

### Materials and Methods

#### Animals

The Animal Care and Use Committee of the University of Toyama, in compliance with the guidelines of the National Institutes of Health, approved the use and care of animals in this study. Five-week-old [for the repeated maximum electroconvulsive shock (rMECS) experiments] and 8-week-old [for the repeated high-frequency tetanic stimulation (rHFS) experiments] Wister ST male rats were purchased from Sankyo Laboratory Japan Inc. Food and water were provided *ad libitum*. All animals were maintained at room temperature and a 12:12 h light-dark cycle (8 am-8 pm).

#### Ibotenic acid (IBO) injection

For the LTP saturation experiments (Fig. 1, 2 and 3) in freely moving animals, rats underwent complete unilateral hippocampal (left hemisphere) lesions by IBO (Santa Cruz Biotechnology) injection at multiple sites (Jarrard, 1989) ( Table S1). The IBO was dissolved in phosphate-buffered saline (PBS) at a concentration of 10 mg ml^−1^ and a pH of 7.4. The dissolved IBO solution was divided into aliquots and stored in a −30°C freezer. On the day of injection, an IBO aliquot was thawed at room temperature and used for a maximum of 2 days, with any remaining solution then being discarded. During the surgery, rats were anesthetized with an intraperitoneal (i.p.) injection of sodium pentobarbital (65 mg kg^−1^ of body weight). To achieve a complete lesion of the left hippocampus, IBO was injected into multiple sites (14 sites; Table S1) using a 2 ml Hamilton syringe and a guide cannula (EIM-362; Eicom, San Diego, CA). The quantity of IBO injected at each site varied from 0.05 to 0.1 μl. The rate of IBO injection was 0.1 μl min^−1^, with the microinjection pump pressure set at 30% (Legato 111, KD Scientific, Holliston, MA). The guide cannula was kept in place for an additional minute to prevent spreading. After completion of injections, wounds were closed with dental cement, and the animals were placed in their home cage (HC) and their health condition was monitored until recovery from anesthesia was complete. After completing the full course of experiments, post-hoc analyzes were performed in all animals to check that the lesion to the left side of the hippocampus was complete; this was performed by staining with cresyl violet.

#### Saturation of hippocampal LTP in freely moving rats

Ten days after the IBO injection into the left-side hippocampus, a second surgery was conducted to implant electrodes into the right-side hippocampus. A bipolar stimulating electrode was positioned in the perforant pathway (PP) to selectively stimulate the PP and projections, while a monopolar recording electrode was placed in the dentate gyrus (DG) (Fig. 1A). Tungsten wire was used to make the electrodes. The stimulating electrode was placed 7.8 ± 0.3 mm posterior, 4.8 ± 0.3 mm lateral and 4.2 ± 0.3 mm inferior to the bregma. The recording electrode was implanted ipsilaterally, at 4.0 ± 0.2 mm posterior, 2.5 ± 0.3 mm lateral and 3.5 ± 0.3 mm inferior to the bregma. Rats were permitted to recover for at least 10 days in individual HCs. Throughout the recovery time, input/output (I/O) curves were determined as a function of current intensity (0.1–1.0 mA). A current that evoked 50% of the maximum fEPSP slope amplitude was used for all rHFS and test pulse experiments, and then the value was kept constant throughout the entire experiment. Test stimuli were delivered at 20 sec intervals to record the fEPSP. LTP was induced by delivering HFS as described previously, with slight modifications (Fukazawa et al., 2003; Matsuo et al., 2000). The HFS (500) was used to stimulate the PP and projections, with each HFS (500) containing 10 trains, with 1 min inter-train intervals. Each train consisted of 5 bursts of 10 pulses at 400 Hz, delivered at 1 sec inter-burst intervals. After baseline recording for 15 min, HFS was induced by a series of six tetanization sessions in 3 days: two tetanizing episodes per day, with a 1.5 h inter-stimulation interval. The seventh tetanization was given on the fourth day to check whether the hippocampal synaptic plasticity was saturated, or whether there would be further cumulative potentiation. Synaptic responses were collected from the DG for 15 min after each HFS session.

#### rMECS stimulation

The rats received a series of 10 repeated MECS, one every 2 days, for 20 days(Stewart et al., 1994). These were given under isoflurane anesthesia (2% isoflurane with 2% O_2_), with the MECS (100 Hz, 0.5 ms, 1 sec, 65 mA) being induced by a pulse generator (UGO BASILE, 57800-001) and the electrodes attached to the ears via an ear clip. After delivering the shock, the ear clip was left in place for an additional 10 sec, before being removed. These parameters were sufficient to induce a seizure in all animals. The sham group received the relevant procedures, except for the shock stimulation.

#### DG-LTP in urethane-anesthetized animals

After the behavioral (contextual fear conditioning; CFC) analysis, rMECS-treated or sham rats were used for the analysis of DG-LTP under urethane anesthesia(Inokuchi et al., 1996), as shown in Fig. 4, C and D. Rats were anesthetized using urethane (i.p., 1.0 g kg^−1^ body weight), and the stimulating and recording electrodes were implanted unilaterally, in the medial perforant pathway (MPP) and DG, respectively (coordinates, recording electrode: AP: 4.0 ± 0.2, ML: 2.5 ± 0.3, Z: 3.5 ± 0.3, stimulating electrode: AP: 7.8 ± 0.3, ML: 4.5 ± 0.3, Z: 4.2 ± 0.3). All the stimuli were conducted using biphasic square wave pulses (200 μs width) with the current intensity set to the level that evoked 50% of the maximum fEPSP slope amplitude. After monitoring a stable basal transmission for at least 15 min, LTP was induced by delivering HFS (100). HFS (100) contained two trains with 1 min inter-train intervals. Each train consisted of 5 bursts of 10 pulses at 400 Hz, delivered at 1 sec inter-burst intervals. The fEPSP slope was monitored by delivering test stimuli at 20 sec intervals. A small animal heat controller (ATC-101B; Unique Medical, Japan) was used to maintain the body temperature of the animals at 37°C throughout the LTP experiments, which were performed under urethane anesthesia.

#### X-ray irradiation

Either 5-week-old rats or IBO-injected rats of approximately 9-11 weeks of age were irradiated. All animals were anesthetized using pentobarbital (i.p. 35 mg kg^−1^ of body weight) before being irradiated. The completely anesthetized animals were placed inside the X-ray irradiation apparatus (MBR-1520R-3, Hitachi, Japan) and irradiated at 150 kV and 5 mA. For irradiation, 0.5 mm aluminum and 0.2 mm copper filters were used. The animal’s entire body was covered with a lead shield to protect it from the X-ray exposure, while the head was left exposed to the irradiation. The distance between the animal’s head skin surface to the irradiation source was 23.3 mm. The procedure lasted for 10 min, delivering 10 Gy at a dose rate of approximately 1 Gy min^−1^. The non-irradiated (0 Gy) group received all the processes except for the irradiation.

#### Running wheel (RW) experiment

Six-week-old rats were placed in cages equipped with a RW for a period of 6 weeks. Cycle computers were attached to the cages with the sensors placed on the back of the wheel to analyze usage. One day after X-ray irradiation, rats were placed inside the cages equipped with a RW. The non-runner group was kept in the HC. Two weeks later, IBO injection and electrode implantation were performed in the left and right hemispheres of the brain. The rats were allowed a 5 day recovery period in the HC, before being returned to the RW cages for the rest of the experiment.

#### Contextual fear conditioning

All the rats were maintained in individual HCs with laboratory bedding. Training and testing sessions were conducted during the daytime in a soundproof behavioral room. The behavioral room was adjacent to the animal living room. The behavioral experiments were performed according to a previous description(Kitamura et al., 2009). The conditioning chamber consisted of a Plexiglas front and black side and back walls (width × depth × height: 175 × 165 × 300 mm). The chamber floor consisted of 21 stainless steel rods with a diameter of 2 mm, placed 4 mm apart. A shock generator was connected to the rods via a cable harness. During the training phase, rats were placed in the conditioning chamber, and after 3 min the animals received a single foot-shock for 2 sec (1.3 mA for HFS-treated animals, Fig. 1 to 3; and 1.6 mA for rMECS-treated animals, Fig. 4). After the shock, animals remained in the chamber for an additional 1 min, before being returned to their HCs. After 1 day, the rats were placed back into the conditioning chamber for 4 min for a test session. After the end of each session, rats were returned to their HCs and the floor of the chamber was cleaned with distilled water and 70% ethanol. Scoring of the freezing response was conducted using an automated video tracking system (Muromachi Kikai). A freezing score was counted after 3.0 sec of sustained freezing.

#### Auditory fear conditioning

Auditory fear conditioning (AFC) was used as a hippocampus-independent learning task. The conditioning chamber was similar to that used in the CFC experiments. During the training session, rats were placed in the conditioning chamber for 3 min. After 2 min, rats received a tone (85 dB, 1 kHz, 20 sec) co-terminated with a foot-shock (the same as in the CFC task). After the shock, rats remained in the chamber for an additional 40 sec, before being returned to their HCs. After 1 day, the rats were placed in a different chamber (triangular in shape) for the test. The test session lasted for 4 min, with a 2 min pre-tone session followed by a continuous tone being delivered for 2 min. After the end of each session, the rats were returned to their HCs and the chamber’s floor was cleaned with distilled water and 70% ethanol. Freezing response was measured using the automated video tracking system.

#### Motility calculation

The motility of the rats during the CFC was measured using the previously mentioned video tracking system (Kitamura et al., 2012; Kitamura et al., 2009). Motility was calculated for the first 3 min of the conditioning session (before receiving the shock) as the area covered by the rat’s movement.

#### Cresyl violet staining

The procedure for cresyl violet staining was similar to that described previously (Deitch and Moses, 1957; Humason, 1983). Animals were deeply anesthetized using an overdose injection of pentobarbital and then perfused transcardially using cold PBS (pH 7.4) followed by 4% paraformaldehyde in PBS. The brains were removed and further post-fixed in 4% paraformaldehyde in PBS at 4°C for at least 72 h. Horizontal sections (20 μm) were cut on a cryostat, and five sections were collected from the hippocampus for staining. Brain sections were inoculated for 12-14 min in the cresyl violet solution, which was kept in an oven (60°C). After washing the sections with distilled water, they were dehydrated with 70% Alcohol, 95% Alcohol, 100% Alcohol and then Xylene.

#### BrdU and NeuN labeling and quantification

Procedures for the quantification of hippocampal cell proliferation and neurogenesis followed those described previously (Kitamura et al., 2009; Ohkawa et al., 2012). For counting of cell proliferation, rats received a single injection of BrdU (100 mg kg^−1^, i.p., Sigma; BrdU dissolved in 0.9% NaCl solution), and then 2 h after the BrdU injection, they were anesthetized using an overdose of pentobarbital solution and transcardially perfused with cold PBS followed by 4% paraformaldehyde in PBS. For analysis of neurogenesis in the rHFS and test pulse groups, rats received three BrdU injections (once per day for 3 successive days) and were perfused 1 day after the CFC experiment. For analysis of neurogenesis in the rMECS and sham groups, rats received six BrdU injections (two per day for 3 consecutive days) and were perfused 1 day after the CFC experiments. After the rats were perfused with paraformaldehyde, the brains were removed and further post-fixed in 4% paraformaldehyde in PBS for 24 h at 4°C. The brains were then equilibrated overnight in 25% sucrose in PBS, before being frozen in dry ice powder. For BrdU staining, 20 μm coronal sections were cut on a cryostat, with one in every three sections from the hippocampus being collected (3.0–5.0 mm from the bregma; 30 sections from each rat). The brain sections were boiled in 0.01 M sodium citrate buffer (pH 6.0) for 10 min and then treated with 2 M HCl for 30 min. The sections were then rinsed with 0.1 M boric acid (pH 8.5) for 10 min. Blocking of the sections was performed using blocking solution (5% donkey serum in PBS containing 0.1% Triton X-100) at room temperature for 1 h. After blocking, sections were incubated with blocking solution containing rat anti-BrdU (1:800, AbD Serotec, Bio-Rad) and mouse anti-NeuN (1:300, Chemicon, EMD Millipore) antibodies. After washing with PBS, the sections were incubated with anti-rat IgG-Alexa Fluor 488 (1:200, Molecular Probes, Invitrogen) and anti-rabbit IgG-Alexa Fluor 546 (1:200, Molecular Probes, Invitrogen) antibodies at room temperature for 3 h. The slides were then washed three times with PBS (10 min per wash). Sections were mounted on glass slides with Pro-Long Gold antifade reagents (Invitrogen). Hippocampal fluorescent images were obtained on a Keyence microscope (BZ-9000, Keyence, Japan). After the images were acquired, the numbers of BrdU^+^ and BrdU^+^/NeuN^+^ cells were counted manually, from 30 sections per rat, using software supplied with the Keyence microscope.

#### Statistics

All data are presented as mean ± SEM. Statistical analyzes were performed using GraphPad software (GraphPad Prism 6). The number of animals is indicated by “*n*” Comparisons between the data of two groups were analyzed using unpaired student’s *t*-tests or paired *t*-tests (within the same group). If the data did not meet the assumptions of the *t*-test, the data were analyzed using the nonparametric Mann-Whitney U-test. Multiple group comparisons were assessed using one-way, two-way or repeated-measures analysis of variance (ANOVA), followed by the appropriate post-hoc test when significant main effects or interactions were detected. The null hypothesis was rejected at the *P* < 0.05 level.

## Author Contributions

J.A., T. Kitamura and K.I. conceived and designed the study. J.A. performed the MECS treatment and drug injection. J.A. and Y.S. conducted animal surgery and electrophysiological experiments. J.A. and T. Kitamura performed the behavioral experiments. J.A. and N.O. performed the histological analysis. J.A. and T. Kondo performed X-ray irradiation experiments. J.A., T. Kitamura, and K.I. prepared the manuscript. K.I. guided and supervised the entire project.

## Acknowledgements

We thank Z. Qi for setting up the LTP experiments, and M. Shehata and M. Nomoto for valuable discussion on behavioral analysis and the manuscript. This work was supported by the Core Research for Evolutional Science and Technology (CREST) program (JPMJCR13W1) of the Japan Science and Technology Agency (JST), JSPS KAKENHI grant number JP23220009, a Grant-in-Aid for Scientific Research on Innovative Areas ‘Memory dynamism’ (JP25115002) from the MEXT, the Mitsubishi Foundation, the Uehara Memorial Foundation, and the Takeda Science Foundation support to K.I. T. Kitamura was funded by Sasagawa Scientific Grants (22-440). We thank Nakamura-sekizenkai (K2012011006) for supporting J.A.

## REFERENCES

Altman, J., and Das, G.D. (1965). Autoradiographic and histological evidence of postnatal hippocampal neurogenesis in rats. J Comp Neurol 124, 319–335.

Buzsaki, G., and Moser, E.I. (2013). Memory, navigation and theta rhythm in the hippocampal-entorhinal system. Nat Neurosci 16, 130–138.

Castro, C.A., Silbert, L.H., McNaughton, B.L., and Barnes, C.A. (1989). Recovery of spatial learning deficits after decay of electrically induced synaptic enhancement in the hippocampus. Nature 342, 545–548.

Conway, M.A. (2009). Episodic memories. Neuropsychologia 47, 2305–2313.

de Vivo, L., Bellesi, M., Marshall, W., Bushong, E.A., Ellisman, M.H., Tononi, G., and Cirelli, C. (2017). Ultrastructural evidence for synaptic scaling across the wake/sleep cycle. Science 355, 507–510.

Deitch, A.D., and Moses, M.J. (1957). The Nissl substance of living and fixed spinal ganglion cells. II. An ultraviolet absorption study. J Biophys Biochem Cytol 3, 449–456.

Deng, W., Aimone, J.B., and Gage, F.H. (2010). New neurons and new memories: how does adult hippocampal neurogenesis affect learning and memory? Nat Rev Neurosci 11, 339–350.

Fukazawa, Y, Saitoh, Y., Ozawa, F., Ohta, Y, Mizuno, K., and Inokuchi, K. (2003). Hippocampal LTP is accompanied by enhanced F-actin content within the dendritic spine that is essential for late LTP maintenance in vivo. Neuron 38, 447–460.

Gould, E., Beylin, A., Tanapat, P, Reeves, A., and Shors, TJ. (1999). Learning enhances adult neurogenesis in the hippocampal formation. Nat Neurosci 2, 260–265.

Hebb, D.O. (1949). The Organization of Behavior (New York Wiley and Sons).

Humason, G.L. (1983). Animal Tissue Techniques (San Francisco: W. H. Freeman and Company).

Inokuchi, K., Kato, A., Hiraia, K., Hishinuma, F., Inoue, M., and Ozawa, F. (1996). Increase in activin beta A mRNA in rat hippocampus during long-term potentiation. FEBS Lett 382, 48–52.

Jarrard, L.E. (1989). On the use of ibotenic acid to lesion selectively different components of the hippocampal formation. Journal of Neuroscience Methods 29, 251–259.

Kitamura, T., and Inokuchi, K. (2014). Role of adult neurogenesis in hippocampal-cortical memory consolidation. Molecular brain 7, 13.

Kitamura, T., Okubo-Suzuki, R., Takashima, N., Murayama, A., Hino, T., Nishizono, H., Kida, S., and Inokuchi, K. (2012). Hippocampal function is not required for the precision of remote place memory. Molecular brain 5, 5.

Kitamura, T., Saitoh, Y., Takashima, N., Murayama, A., Niibori, Y, Ageta, H., Sekiguchi, M., Sugiyama, H., and Inokuchi, K. (2009). Adult neurogenesis modulates the hippocampus-dependent period of associative fear memory. Cell 139, 814–827.

Li, W., Ma, L., Yang, G., and Gan, W.B. (2017). REM sleep selectively prunes and maintains new synapses in development and learning. Nat Neurosci 20, 427–437.

Lledo, P.M., Alonso, M., and Grubb, M.S. (2006). Adult neurogenesis and functional plasticity in neuronal circuits. Nat Rev Neurosci 7, 179–193.

Matsuo, R., Murayama, A., Saitoh, Y., Sakaki, Y., and Inokuchi, K. (2000). Identification and cataloging of genes induced by long-lasting long-term potentiation in awake rats. J Neurochem 74, 2239–2249.

Moser, E.I., Krobert, K.A., Moser, M.B., and Morris, R.G. (1998). Impaired spatial learning after saturation of long-term potentiation. Science 281, 2038–2042.

Nabavi, S., Fox, R., Proulx, C.D., Lin, J.Y., Tsien, R.Y., and Malinow, R. (2014). Engineering a memory with LTD and LTP. Nature 511, 348–352.

Ofen, N., Kao, Y.C., Sokol-Hessner, P., Kim, H., Whitfield-Gabrieli, S., and Gabrieli, J.D. (2007). Development of the declarative memory system in the human brain. Nat Neurosci 10, 1198–1205.

Ohkawa, N., Saitoh, Y., Tokunaga, E., Nihonmatsu, I., Ozawa, F., Murayama, A., Shibata, F., Kitamura, T., and Inokuchi, K. (2012). Spine formation pattern of adult-born neurons is differentially modulated by the induction timing and location of hippocampal plasticity. PLoS One 7, e45270.

Scoville, W.B., and Milner, B. (1957). Loss of recent memory after bilateral hippocampal lesions. J Neurol Neurosurg Psychiatry 20, 11–21.

Stewart, C., Jeffery, K., and Reid, I. (1994). LTP-like synaptic efficacy changes following electroconvulsive stimulation. Neuroreport 5, 1041–1044.

Toni, N., Teng, E.M., Bushong, E.A., Aimone, J.B., Zhao, C., Consiglio, A., van Praag, H., Martone, M.E., Ellisman, M.H., and Gage, F.H. (2007). Synapse formation on neurons born in the adult hippocampus. Nat Neurosci 10, 727–734.

Tononi, G., and Cirelli, C. (2014). Sleep and the price of plasticity: from synaptic and cellular homeostasis to memory consolidation and integration. Neuron 81, 12–34.

van Praag, H., Kempermann, G., and Gage, F.H. (1999). Running increases cell proliferation and neurogenesis in the adult mouse dentate gyrus. Nat Neurosci 2, 266–270.

van Praag, H., Shubert, T., Zhao, C., and Gage, F.H. (2005). Exercise enhances learning and hippocampal neurogenesis in aged mice. J Neurosci 25, 8680–8685.

Villeda, S.A., Luo, J., Mosher, K.I., Zou, B., Britschgi, M., Bieri, G., Stan, T.M., Fainberg, N., Ding, Z., Eggel, A., et al. (2011). The ageing systemic milieu negatively regulates neurogenesis and cognitive function. Nature 477, 90–94.

Whitlock, J.R., Heynen, A.J., Shuler, M.G., and Bear, M.F. (2006). Learning induces long-term potentiation in the hippocampus. Science 313, 1093–1097.

Zhao, C., Deng, W., and Gage, F.H. (2008). Mechanisms and functional implications of adult neurogenesis. Cell 132, 645–660.

